# Signal triangulation coordinates cell fate decisions in the developing jaw

**DOI:** 10.64898/2026.04.07.716963

**Authors:** Eric Paulissen, Muniza Junaid, Lyla Brugger, Hung-Jhen Chen, J. Gage Crump

**Affiliations:** Eli and Edythe Broad Center for Regenerative Medicine and Stem Cell Research, Department of Stem Cell Biology and Regenerative Medicine, Keck School of Medicine, University of Southern California, Los Angeles, CA 90033, USA

## Abstract

Development of the vertebrate lower jaw depends on spatially precise cell fate decisions by cranial neural crest-derived cells (CNCCs) of the mandibular arch. From closer to the mouth (oral) to farther away (aboral), CNCCs adopt bone, cartilage, tendon, and stromal fates to shape the jaw skeleton and ensure proper connectivity to the musculature. How signaling pathways impact downstream transcriptional programs to generate distinct cell fates along the oral-aboral axis remains incompletely understood. Using photoconversion-based lineage tracing of CNCCs in zebrafish, we show that oral cells contribute to the lower jaw skeleton and aboral cells primarily to tendon, ligament, and stromal tissues. During embryogenesis, the oral domain is partitioned into lateral *pitx1*+ and medial *foxf1*+ subdomains distinct from an aboral *nr5a2*+, *gsc*+ domain. Using pharmacological inhibition and transgenic misexpression, we find that Bmp signaling establishes aboral *nr5a2* and *gsc* expression, Fgf signaling oral-lateral *pitx1* expression, and Hedgehog signaling oral-medial *foxf1* expression. Analysis of mutants for *pitx1*, *nr5a2*, and *gsc* reveal that their oral-aboral expression domains are established independently of each other. We also identify enhancers regulating oral-aboral expression of *pitx1* and *nr5a2*, with mutagenesis showing roles for Fgf-dependent ETS motifs in oral *pitx1* and Bmp-dependent E-box motifs in aboral *nr5a2* enhancer activity, consistent with dependence of *nr5a2* expression on the Bmp-dependent E-box factor Hand2. These findings reveal how triangulation of three major signaling pathways converge on distinct gene regulatory modules to establish distinct oral-aboral cell fate decisions in the developing jaw.

## INTRODUCTION

The connective tissues and skeletal elements of the jaw arise from CNCCs organized into a series of segments called pharyngeal arches (Platt, 1893). CNCCs of the first (mandibular) arch give rise to Meckel’s cartilage and associated bones of the jaw, as well as tendons and ligaments, salivary glands, and other mesenchymal derivatives (Baker et al., 1997; Evans and Noden, 2006; Nakamura, 1982; Noden, 1978). Prior to differentiation, arch CNCCs are organized along multiple regional axes that in part prefigure their eventual fate. For example, the lack of Homeobox (Hox) transcription factor expression promotes a jaw fate of mandibular arch CNCCs compared to more posterior hyoid and branchial arches (Gendron-Maguire et al., 1993; Miller et al., 2004; Rijli et al., 1993). Along the dorsoventral axis of zebrafish (equivalent to the proximodistal axis in mouse), the Pou3f3 transcription factor promotes a dorsal/proximal upper jaw fate, Distal-less (Dlx) 5/6 transcription factors promote a ventral/distal lower jaw fate, and the Hand2 E-box transcription factor promotes the most distal portion of the lower jaw (Barske et al., 2020; Beverdam et al., 2002; Jeong et al., 2008; Miller et al., 2003; Yanagisawa et al., 2003). These domains are established in part through Jagged-Notch signaling dorsally, and a combination of Bmp and Endothelin 1 (Edn1) signaling ventrally (Alexander et al., 2011; Talbot et al., 2010; Zuniga et al., 2010; Zuniga et al., 2011).

Compared to anteroposterior and dorsoventral patterning, less is known about how the oral versus aboral axis of the mandibular arch is established. In most vertebrates, including many fishes, the oral domain gives rise to teeth, with progressively more aboral CNCCs giving rise to first jaw skeletal tissues and then tendons, ligaments, and salivary gland mesenchyme (Chen et al., 2023; Fraser et al., 2009; Soukup et al., 2008). Zebrafish, however, lack oral teeth, making it unclear whether oral-aboral mandibular arch patterning would be the same as in other vertebrates. One conserved marker of the oral mandibular domain in mouse and zebrafish is *Pitx1/pitx1*, with *Gsc/gsc* marking the aboral mandibular domain as well as other mandibular and hyoid arch regions (Mitsiadis and Drouin, 2008; Schulte-Merker et al., 1994; Yamada et al., 1995). Recently, we identified *Nr5a2/nr5a2* as another conserved marker of the aboral mandibular domain in mice and fish (Chen et al., 2023). Consistent with distinct roles in oral versus aboral patterning, the lower jaw skeleton and mandibular teeth are reduced in mouse *Pitx1* mutants (Mitsiadis and Drouin, 2008; Szeto et al., 1999), whereas lower jaw tendons and salivary glands are reduced and missing, respectively, in CNCC-specific mouse *Nr5a2* mutants, with jaw tendons similarly lost in zebrafish *nr5a2* mutants (Chen et al., 2023).

Prior work in mouse and zebrafish has shown that mandibular *Pitx1* and *Gsc* expression requires Edn1 signaling and its target gene Hand2 (Askary et al., 2017; Clouthier et al., 1998), with ectopic *Edn1* or *Hand2* expression resulting in duplicated *Gsc/Pitx1* expression in the maxillary domain (Funato et al., 2016; Sato et al., 2008). In cultured mouse mandibular explants, Bmp4 beads inhibit *Pitx1* expression, with Fgf8 beads reciprocally inducing *Pitx1* and inhibiting *Gsc* expression (Tucker et al., 1999). However, ectopic Bmp activity also results in loss of oral ectodermal *Fgf8* in chick embryos (Shigetani et al., 2000), suggesting that the inhibitory role of Bmp on *Pitx1* expression could be indirect due to loss of Fgf signaling. Bmp signaling also induces *hand2* expression, with *hand2* function required for *gsc* expression (Alexander et al., 2011; Talbot et al., 2010; Zuniga et al., 2011). These experiments suggest that Fgf8 from the oral ectoderm and Bmp4 from the ventral ectoderm could interact to pattern the oral-aboral axis, though their mechanisms of action remain incompletely understood.

In addition to Fgf8 and Bmp4, the Sonic Hedgehog (Shh) ligand is produced by the underlying medial endoderm, which in turn induces Shh in the oral ectoderm (Balczerski et al., 2012; Brito et al., 2006). CNCC-specific loss of Hedgehog (Hh) signaling results in expanded Bmp activity, a smaller mandible, and development of ectopic bone in the oral domain (Jeong et al., 2004; Schwend and Ahlgren, 2009; Xu et al., 2019a). Mandibular Hh signaling functions in part by inducing expression of *Foxf1/foxf1* and *Foxf2/foxf2*, with CNCC-specific loss of *Foxf1/Foxf2* in mouse resulting in similar ectopic bone as seen upon loss of Hh activity (Xu et al., 2018; Xu et al., 2019a). In zebrafish, mutation of *foxf1/foxf2* orthologs results in reductions of lower jaw Meckel’s cartilage, with misexpression of *foxf1* resulting in a loss of dermal bone and expansion of jaw cartilage (Xu et al., 2018). Hh signaling and Foxf1/2 therefore appear to have a role in promoting cartilage medially in the mandibular arch, distinct from the role of Fgf8 and Pitx1 in oral bone and tooth formation and from Nr5a2 in aboral tendon and gland formation. How Hh, Fgf, and Bmp signaling interact to establish gene expression domains prefiguring distinct cell fates in the developing jaw has remained unclear.

In this study, we combined pharmacological, mutant, and transgenic manipulation of signaling activity with enhancer testing to reveal how distinct oral-aboral domains are established in zebrafish. We found that Bmp signaling establishes an *nr5a2*+; *gsc*+ aboral domain, with Fgf and Hh signaling inducing partially overlapping *pitx1*+ lateral and *foxf1*+ medial portions, respectively, of the oral domain. By transgenic testing of putative enhancers from single-nuclei assay for transposase accessible chromatin and sequencing (snATACseq) datasets for zebrafish CNCCs (Fabian et al., 2022), we identified oral *pitx1* and aboral *nr5a2* enhancers, with the *pitx1* enhancer being sequence-conserved in mammals. We further showed requirement of ETS sites for oral *pitx1* enhancer activity, likely reflecting Fgf regulation of multiple ETS genes (Roehl and Nüsslein-Volhard, 2001), and requirement of E-box sites for aboral *nr5a2* enhancer activity consistent with regulation by the Bmp-dependent E-box factor Hand2 (Alexander et al., 2011; Zuniga et al., 2011). These findings illuminate how Fgf, Hh, and Bmp signaling establish spatially distinct downstream transcription factor programs to regulate the specification of distinct skeletal and connective tissue cell types along the mandibular oral-aboral axis.

## RESULTS

### Fate map of mandibular arch CNCCs along the oral-aboral axis

To understand the types of cells arising from distinct domains along the mandibular oral-aboral axis of zebrafish, we performed photoconversion-based lineage tracing using a CNCC-enriched *sox10:kikGR* transgenic line. We used a UV laser to photoconvert kikGR from green to red in 3 distinct domains of the ventral mandibular arch at 36 hours post-fertilization (hpf), and then re-imaged at 4 days post-fertilization (dpf) to assess contributions to CNCC derivatives (**Figure 1A-C**). We took advantage of the fact that *sox10:kikGR* has a second wave of expression in chondrocytes, resulting in the jaw cartilages being labeled green as a reference. Photoconversion of the domain closest to the oral ectoderm resulted in labeling of mesenchyme prefiguring the dentary bone and distal Meckel’s cartilage. Whereas photoconversion of an oral-aboral intermediate domain resulted in labeling of proximal Meckel’s cartilage, tendon, and some ligament cells, photoconversion of the most aboral domain resulted in labeling of only ligament and stromal cells posterior to Meckel’s cartilage. These results are consistent with the zebrafish oral mandibular domain giving rise to bone and cartilage, and the aboral domain giving rise to soft connective tissues and stromal cells (**Figure 1D)**. As confocal illumination uses a beam of light, one caveat is that we labeled CNCCs throughout the mediolateral axis in each domain and thus could not discriminate whether cartilage and other fates arose preferentially from medial or lateral domains.

**Figure 1.**
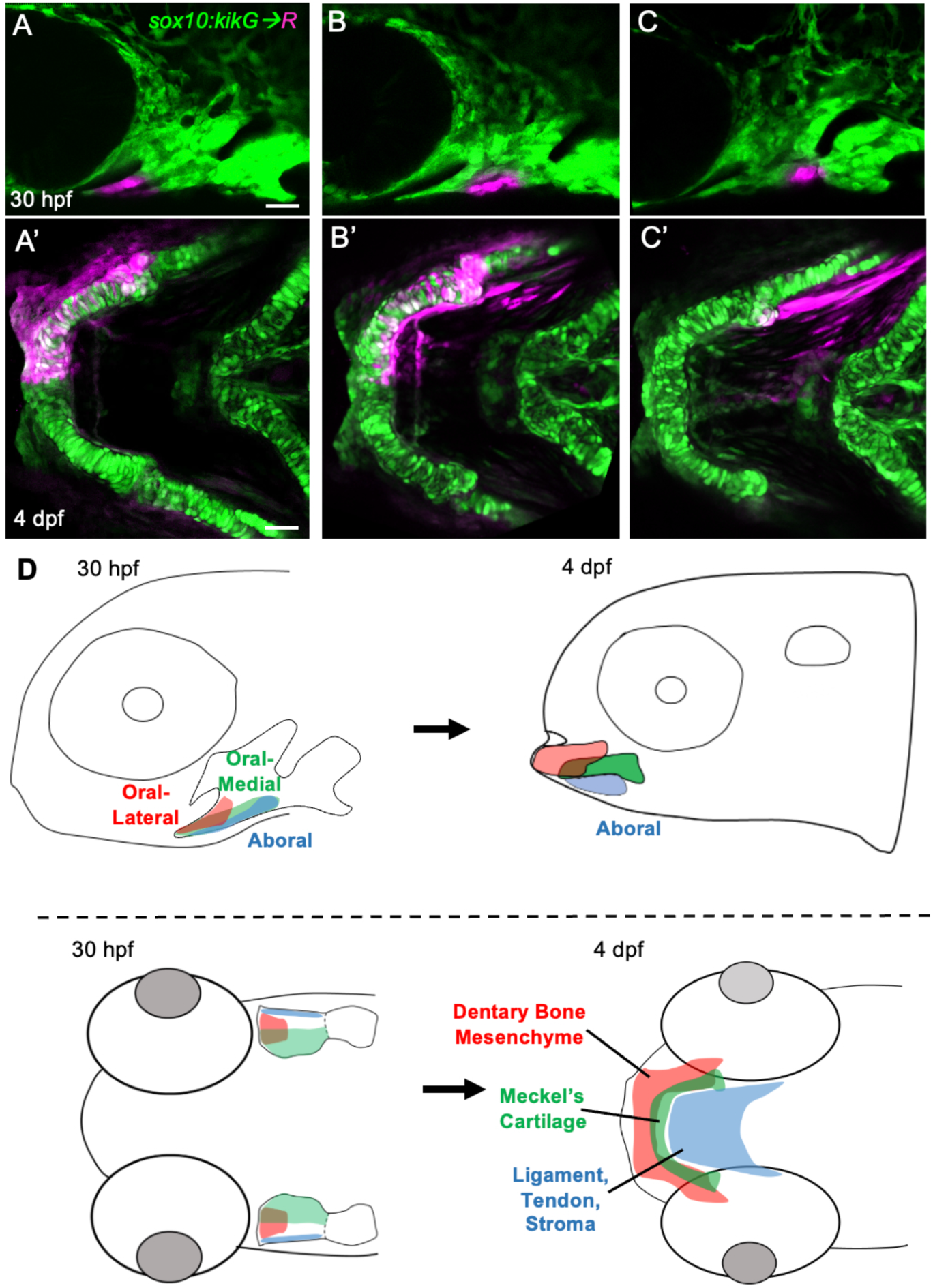
Lineage tracing of the oral-aboral axis of the ventral mandibular domain. **(A-C)** *sox10:kikGR* labels CNCCs of the first two pharyngeal arches in lateral view at 30 hpf and facial cartilages in ventral view at 4 dpf. The unconverted kikGR protein is green. UV light was used for photoconversion of kikGR to red (pseudocolored magenta) in the oral (A), oral-aboral intermediate (B), or aboral (C) domains at 30 hpf, with re-imaging of the same animals at 4 dpf revealing differential contributions to dentary bone-associated oral mesenchyme, Meckel’s chondrocytes, and ligament, tendon, and stromal cells. Scale bars = 25 μM. (**D**) Schematics in lateral (top) and ventral (bottom) views of distinct domain contributions along the oral-aboral axis.

### Gene expression analysis reveals three distinct domains in the ventral mandibular arch that correlate with Bmp, Fgf, or Hh activity

We next sought to determine how oral-aboral fate domains correspond to embryonic gene expression in zebrafish. To do so, we performed multiplexed in situ RNA hybridization chain reaction (HCR) at 36 hpf in combination with *sox10:GFPCAAX* to highlight CNCCs (detected by anti-GFP antibody) (**Figure 2A-K**). Whereas *pitx1* was expressed in CNCCs along the oral ectoderm, *nr5a2* and *gsc* expression co-localized in aboral domain CNCCs along the ventral mandibular arch margin. Both oral *pitx1* and aboral *nr5a2* domains co-localized with *hand2* expression, although *pitx1* expression also extended more dorsally into a *hand2*-negative region. Expression of *foxf1* partially overlapped with *pitx1*, with a *foxf1-*only domain extending medially and a *pitx1*-only domain extending laterally. When viewed along the mediolateral axis, aboral genes *nr5a2* and *gsc* were also expressed in more lateral positions than *foxf1* and *pitx1*. Of note, *gsc* has additional arch expression in the ventral hyoid and dorsal mandibular and hyoid domains, as well as in the medial-most regions of the maxillary and mandibular domains. These distinct expression domains also match predictions from our integrated scRNAseq and snATACseq datasets of zebrafish CNCCs (Fabian et al., 2022), with additional predicted aboral-restricted genes shown in **Figure S1** and **Table S1**. Our analysis therefore identifies three distinct domains within the ventral mandibular arch labeled by *pitx1* (oral-lateral), *foxf1* (oral-medial), and *nr5a2* and *gsc* (aboral).

**Figure 2.**
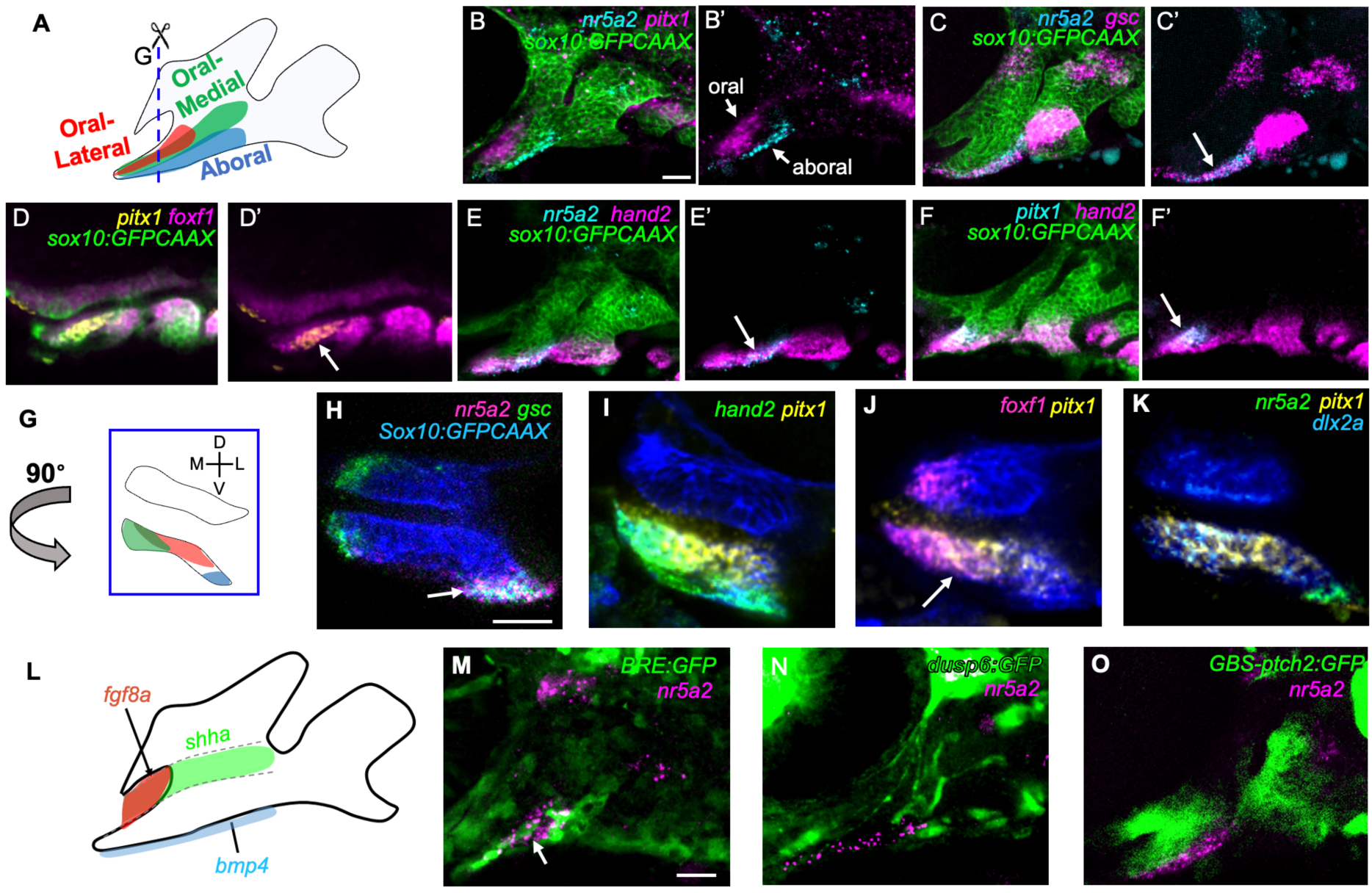
Oral-aboral gene expression relative to signaling pathway activity. **(A)** Schematic of the first two pharyngeal arches in lateral view with mandibular sub-domains colored. **(B-F)** Combinatorial in situ hybridizations for the indicated genes at 36 hpf shown in lateral view. All images are projections through the arches, except for (D) which is a partial projection through the medial aspects of the arches. Anti-GFP antibody labeling shows *sox10:GFPCAAX*+ CNCCs for reference. White arrows indicate regions of overlapping expression. **(G)** Schematic in coronal view corresponding to the cut site in (A). **(H-K)** Combinatorial in situ hybridizations for the indicated genes at 36 hpf shown in coronal view, with *sox10:GFPCAAX* labeling of CNCCs in blue for reference. Arrows show co-localization of *nr5a2* and *gsc* in lateral, aboral CNCCs (H) and *foxf1* and *pitx1* in mediolaterally intermediate, oral CNCCs (J). **(L)** Schematic of first two pharyngeal arches shows expression of *fgf8a* in the oral ectoderm, *shha* in the medial endoderm (dashed lines), and *bmp4* in the ventral ectodermal margin. **(M-O)** Anti-GFP labeling of the first two pharyngeal arches at 36 hpf in lateral view shows signaling activity for Bmp (*BRE:GFP*), Hh (*GBS-ptch2:EGFP*), and Fgf (*dusp6:GFP*) pathways relative to aboral *nr5a2* expression. Arrow denotes co-localization of *nr5a2* expression and Bmp activity. Scale bars = 25μM.

To identify candidate regulators of oral-aboral domains, we next analyzed the activity of transgenic reporters for Bmp, Fgf, and Hh signaling (**Figure 2L-O**). We found preferential activity of a Bmp-responsive *BRE:GFP* transgene in the *nr5a2*+ aboral domain, and activity of Fgf-responsive *dusp6:GFP* and Hh-responsive *GBS-ptch2:GFP* transgenes in *nr5a2*- oral-lateral and oral-medial domains, respectively. These findings point to Bmp, Fgf, and Hh signaling as candidate regulators of oral-aboral expression domains in zebrafish.

### Distinct roles for Bmp, Fgf, and Hh signaling in oral-aboral patterning

To analyze roles for Bmp signaling in oral-aboral gene expression, we treated embryos with the Bmp receptor antagonist DMH1 shortly after pharyngeal arch formation (from 18-36 or 22-36 hpf). Consistent with co-localization of Bmp reporter activity with *nr5a2* expression, DMH1-mediated Bmp inhibition resulted in a loss of *nr5a2* and *gsc* expression specifically in the aboral domain of the ventral mandibular arch (**Figure 3A,B**). A similar loss of aboral *nr5a2* expression was seen using Dorsomorphin to inhibit Bmp signaling (**Figure 3G,H**), and we confirmed effective DMH1 inhibition of Bmp signaling by loss of the known Bmp target gene *hand2* (**Figure 3D**). In contrast, oral *pitx1* expression was unaffected in DMH1-treated embryos, and medial *foxf1* expression was only mildly reduced (**Figure 3C-F**). Reciprocally, embryo-wide misexpression of Bmp4 by heat shock of *UAS:Bmp4; hsp70l:Gal4* embryos at 22 hpf resulted in expansion of *gsc* and *nr5a2* expression throughout the mandibular arch and loss of *pitx1* oral expression (**Figure 3I-L**).

**Figure 3.**
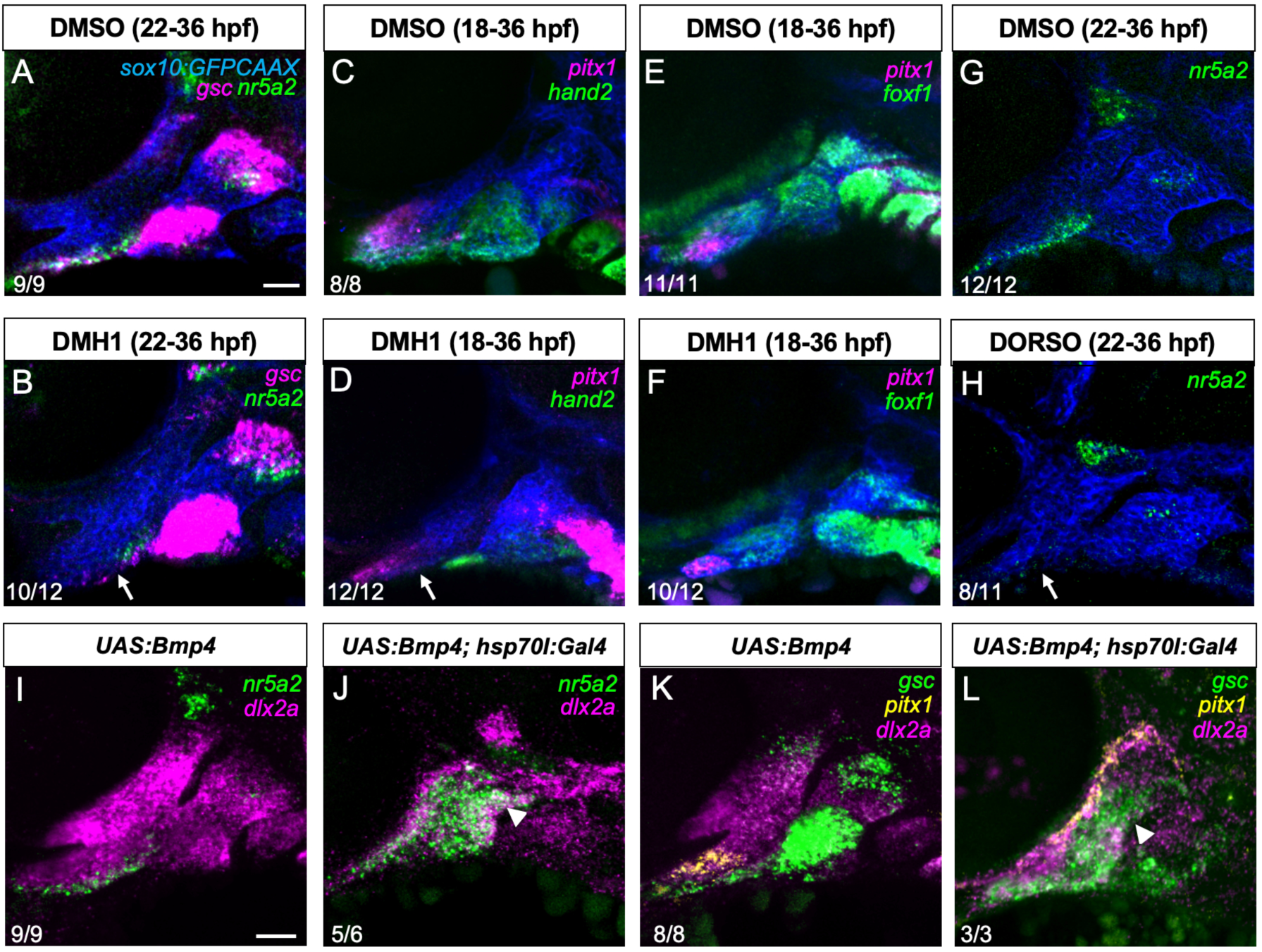
BMP promotes aboral gene expression. **(A-H)** Combinatorial in situ hybridizations for the indicated genes at 36 hpf shown in lateral view, with anti-GFP antibody showing *sox10:GFPCAAX*+ CNCCs of the first two pharyngeal arches for reference. Compared to DMSO-treated controls (A-D), embryos treated with the Bmp antagonists DMH1 and Dorsomorphin (DORSO) for the indicated time intervals show loss of aboral mandibular *gsc* and *nr5a2* and ventral *hand2* expression (white arrows) but not oral *pitx1*, medial *foxf1*, or second arch *gsc* expression. Proportions of embryos with the shown patterns are listed. **(I-L)** Combinatorial in situ hybridizations for the indicated genes at 36 hpf shown in lateral view, with *dlx2a* labeling CNCCs of the first two pharyngeal arches for reference. Compared to *UAS:Bmp4* controls, *hsp70l:Gal4; UAS:Bmp4* embryos subjected to heat-shock at 22 hpf to induce embryo-wide Bmp4 misexpression showed expansion of *n5a2* and *gsc* expression throughout the mandibular arch and loss of *pitx1* expression in the oral domain (white arrowhead indicates expanded areas). Proportions of embryos with the shown patterns are listed. Scale bars = 25 μM.

We next examined roles of Fgf and Hh signaling using a similar pharmacological approach. Inhibition of Fgf signaling with SU5402 from 22 to 36 hpf resulted in an expansion of aboral *nr5a2* and *gsc* gene expression into oral and medial regions of the mandibular arch and loss of oral *pitx1* expression, yet medial *foxf1* expression was largely unaffected (**Figure 4**). In contrast, inhibition of Hh signaling with cyclopamine from 18 to 36 hpf severely reduced *foxf1* expression, consistent with previous reports (Xu et al., 2018; Xu et al., 2019a), but did not affect *pitx1* and *nr5a2* expression (**Figure 5**). These findings support distinct roles for Bmp signaling in establishing the aboral domain, Fgf signaling in establishing the oral-lateral domain, and Hh signaling in establishing the oral-medial domain.

**Figure 4.**
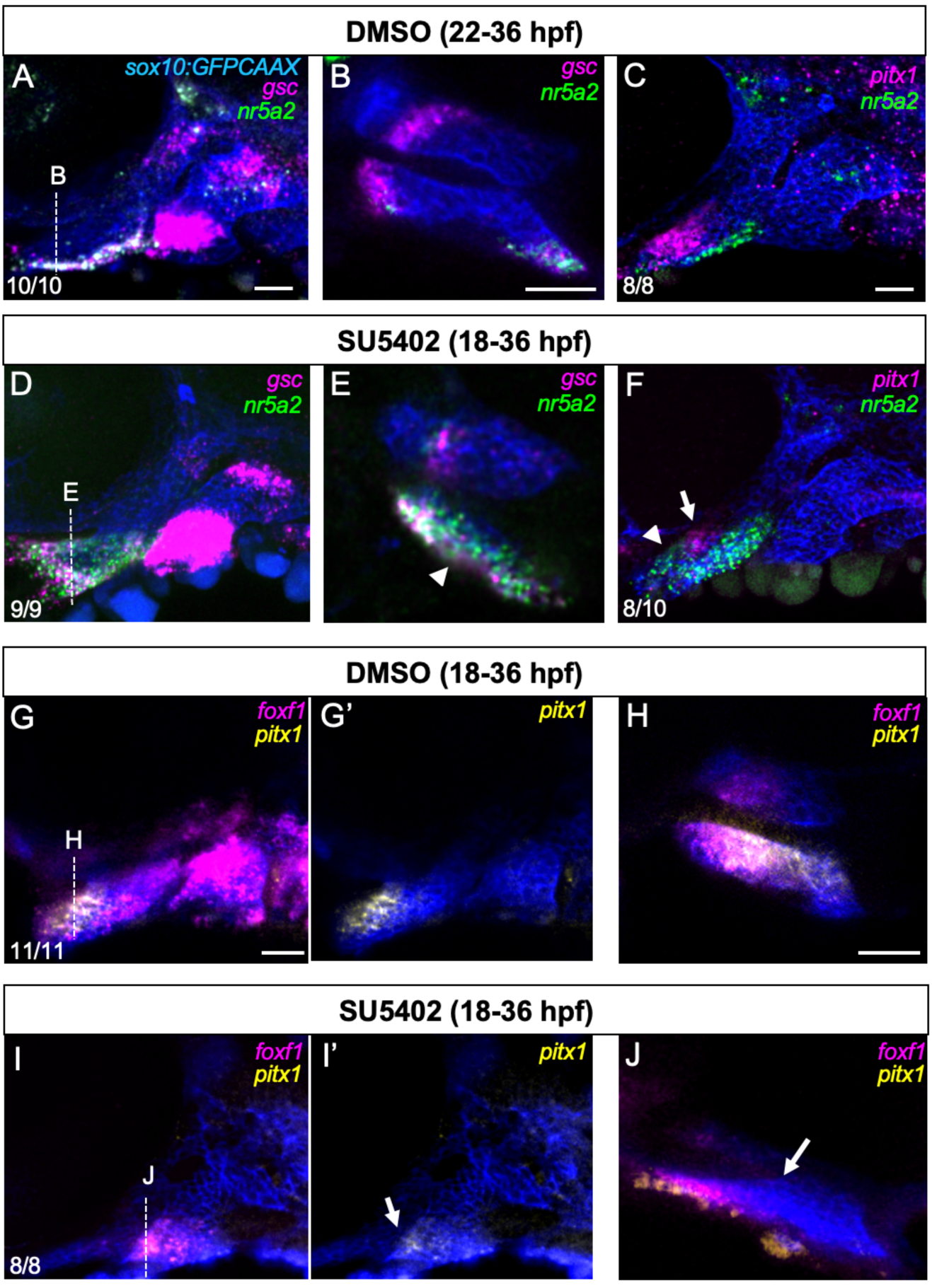
Requirement for FGF signaling in oral gene expression. **(A-J)** Combinatorial in situ hybridizations for the indicated genes at 36 hpf shown in lateral view (A,C,D,F,G,I) and coronal view (B,E,H,J), with anti-GFP antibody showing *sox10:GFPCAAX*+ CNCCs of the first two pharyngeal arches for reference. Compared to DMSO-treated controls, embryos treated with the Fgf antagonist SU5402 for the indicated time intervals showed an expansion of *gsc* and *nr5a2* expression (white arrowheads) into medial (E) and oral (F) domains, and a corresponding severe reduction of *pitx1* expression (I’, J, white arrows). Medial expression of *foxf1* was only moderately affected in the abnormally shaped mandibular arch. Proportions of embryos with the shown patterns are listed. Scale bars = 25 μM.

**Figure 5.**
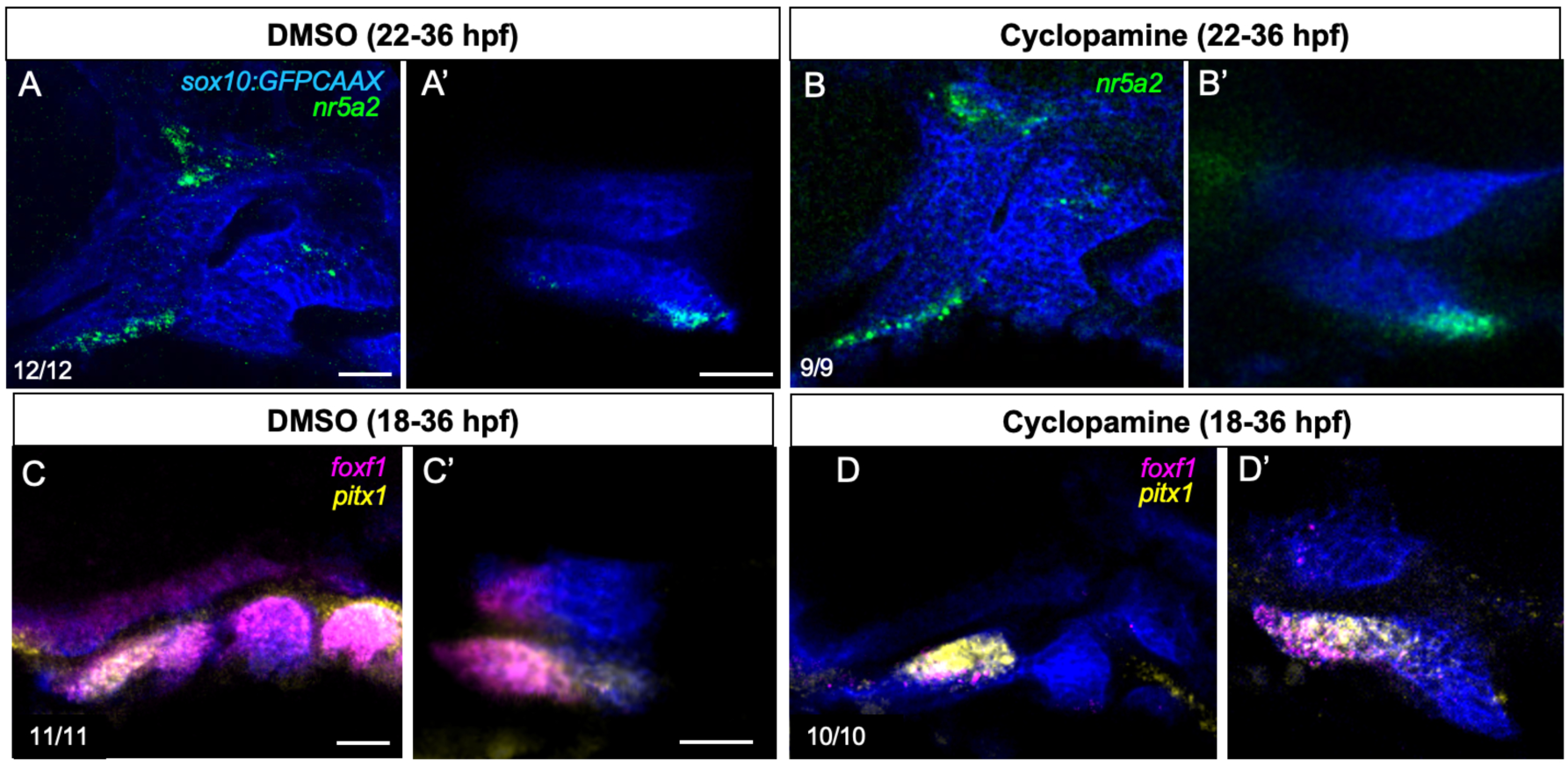
Requirement for Hh signaling in medial gene expression. **(A-D)** Combinatorial in situ hybridizations for the indicated genes at 36 hpf shown in lateral view (A-D) and coronal view (A’-D’), with anti-GFP antibody showing *sox10:GFPCAAX*+ CNCCs of the first two pharyngeal arches for reference. Compared to DMSO-treated controls, embryos treated with the Hh antagonist Cyclopamine for the indicated time intervals show a strong reduction of medial *foxf1* but not aboral *nr5a2* or oral *pitx1* expression. Proportions of embryos with the shown patterns are listed. Scale bars = 25 μM.

### Transcriptional regulation of oral-aboral patterning

We next examined which downstream transcription factors might mediate oral-aboral patterning. We had previously shown that *pitx1* and *foxf1* expression requires Edn1 signaling in zebrafish (Askary et al., 2017), and we find here that *nr5a2* and *gsc* expression are also lost in *edn1* mutants (**Figure 6C,F**). As *hand2* is co-expressed with *nr5a2* and *gsc* and similarly requires Edn1 and Bmp signaling for its expression (Miller et al., 2004; Miller et al., 2007; Zuniga et al., 2011), we examined requirements for Hand2 and found that aboral *nr5a2* and *gsc* but not oral *pitx1* expression is lost in *hand2* mutants (**Figure 6B,E,H**). The positive effects of Bmp signaling on *nr5a2* and *gsc* expression are at least partially dependent on Hand2 function, as *hand2* loss abrogated the ectopic mandibular arch expression of *nr5a2* and *gsc* seen upon embryo-wide Bmp4 misexpression (**Figure 6L-S**). Similar to *Pitx1* regulation in mouse (Bobola et al., 2003), we also found that Hox2 genes likely restrict *nr5a2* expression to the mandibular arch as *nr5a2* expression expanded into the ventral hyoid arch in *moz* mutants that lack *hoxa2b* and *hoxb2a* arch expression (Miller et al., 2004) (**Figure 6I**). In addition, we found that Nr5a2, Gsc, and Pitx1 do not regulate the expression of each other, as *nr5a2* expression was unaffected in *gsc* and *pitx1* mutants (**Figure 6J,K**), and we previously reported normal expression of *gsc* and *pitx1* in *nr5a2* mutants (Chen et al., 2023). Thus, aboral expression of *nr5a2* and *gsc* selectively requires Hand2 function downstream of Bmp and Edn1 signaling but is not negatively regulated by oral Pitx1 or Foxf1 function.

**Figure 6.**
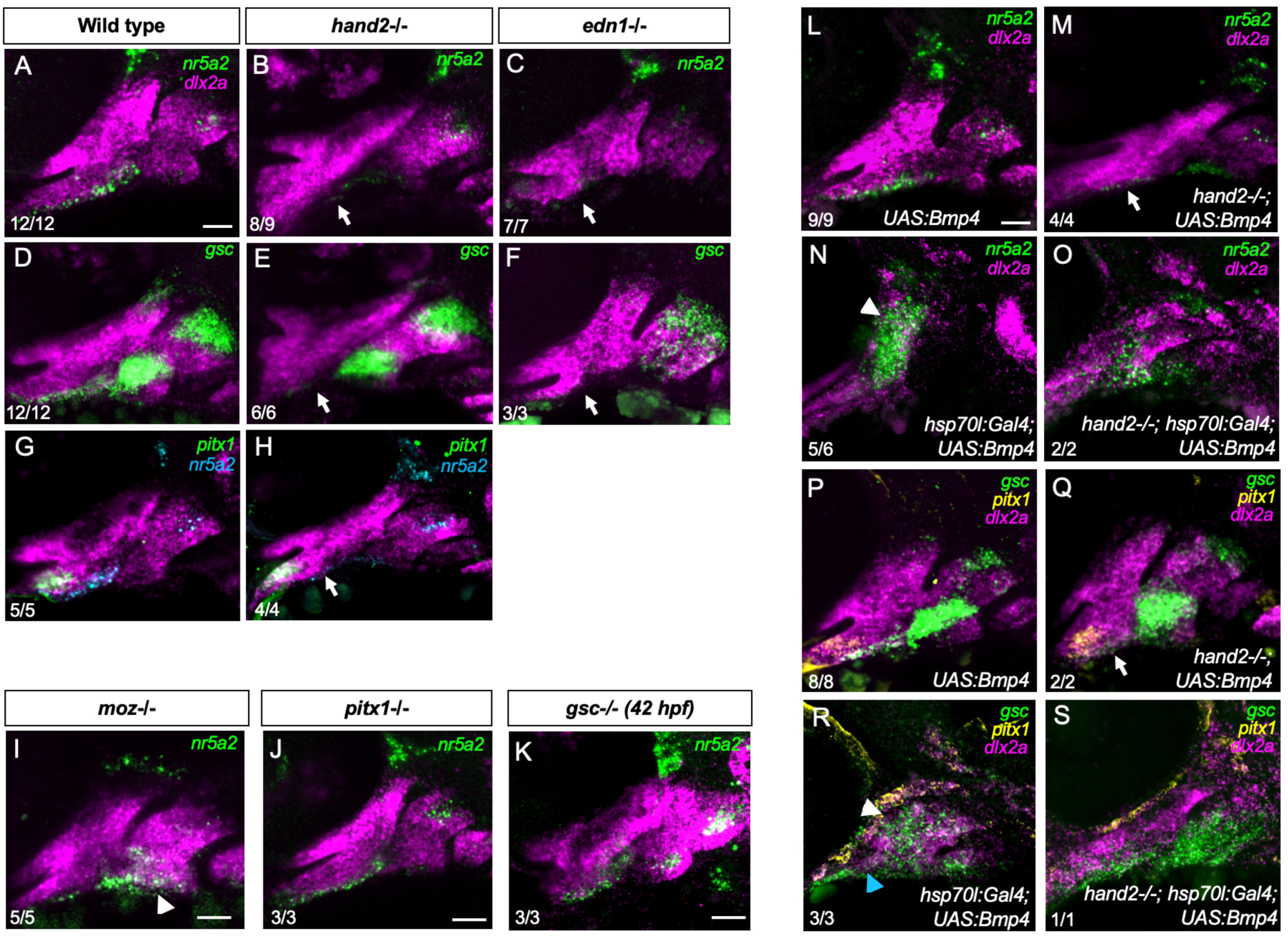
Genetic regulation of oral-aboral gene expression. **(A-K)** In situ hybridizations for the indicated genes at 36 hpf (A-J) or 42 hpf (K) with *dlx2a* expression (magenta) labeling CNCCs of the first two pharyngeal arches for reference. Aboral *nr5a2* and *gsc* expression is lost in *hand2* and *edn1* mutants (arrows), *nr5a2* expression is unaffected in *pitx1* and *gsc* mutants, *pitx1* expression is unaffected in *hand2* mutants, and *nr5a2* expression is expanded into the second arch in *moz* mutants (white arrowhead). **(L-S)** In situ hybridizations for the indicated genes at 36 hpf with *dlx2a* labeling CNCCs for reference. Compared to *UAS:Bmp4* controls, *hand2-/-*; *UAS:Bmp4* embryos lack aboral *nr5a2* and *gsc* but not oral *pitx1* expression (M, Q, arrows), and *hsp70l:Gal4; UAS:Bmp4* embryos subjected to heat-shock at 22 hpf to induce embryo-wide Bmp4 misexpression have expanded *nr5a2* and *gsc* mandibular expression (N, R, white arrowheads) and loss of oral *pitx1* expression (blue arrowhead). Loss of *hand2* in heat-shocked *hand2-/-*; *hsp70l:Gal4; UAS:Bmp4* embryos with Bmp4 misexpression abrogates ectopic *nr5a2* and *gsc* expression in the first arch but does not restore oral *pitx1* expression. Proportions of embryos with the shown patterns are listed. Scale bars = 25 μM.

### Characterization of aboral *nr5a2* and oral *pitx1* enhancers

To further understand the regulatory logic of gene expression along the oral-aboral axis, we examined our integrated scRNAseq and snATACseq datasets of zebrafish CNCCs (Fabian et al., 2022) for differentially accessible chromatin regions (DARs, i.e. putative enhancers) in close proximity to oral-aboral genes at 36 hpf. We identified a DAR ∼111 kb downstream of the *nr5a2* promoter that was selectively accessible in the ventral arches, with accessibility overlapping with *nr5a2* mRNA expression in the aboral mandibular domain and largely non-overlapping with *pitx1* mRNA (**Figure 7A**, **Figure S2**). We also identified a DAR ∼16 kb upstream of the *pitx1* promoter with accessibility overlapping oral *pitx1* mRNA expression and largely excluded from the aboral *nr5a2* mRNA expression domain (**Figure 7B, Figure S2**). In stable transgenic lines, we found activity of *- 111nr5a2:GFP* in the aboral mandibular arch starting around 40 hpf, with GFP expression marking connective tissues posterior to *sox10:DsRed*+ Meckel’s cartilage at 3 and 5 dpf, similar to that of endogenous *nr5a2* as reflected by a *nr5a2:GFPCAAX* knock-in reporter (except for *-111nr5a2:GFP* lacking muscle activity) (**Figure 7C,D**). In contrast, *+16pitx1:nlsEOS* was expressed in the oral domain starting at 40 hpf, persisting in Meckel’s cartilage, the *sp7:BFP*+ dentary bone, and mesenchyme anterior to the cartilage at 3 and 5 dpf (**Figure 7E-G**). At 40 hpf, we confirmed mutually exclusive activity of *-111nr5a2:GFP* and *+16pitx1:nlsEOS* in aboral and oral mandibular domains, respectively (**Figure 7H**). The activities of *+16pitx1:nlsEOS* and *-111nr5a2:GFP* are also largely consistent with the contributions of the oral and aboral mandibular domains, respectively, in our lineage tracing experiments (**Figure 1**), showing that these enhancer activities accurately report the endogenous mandibular expression of these two genes.

**Figure 7.**
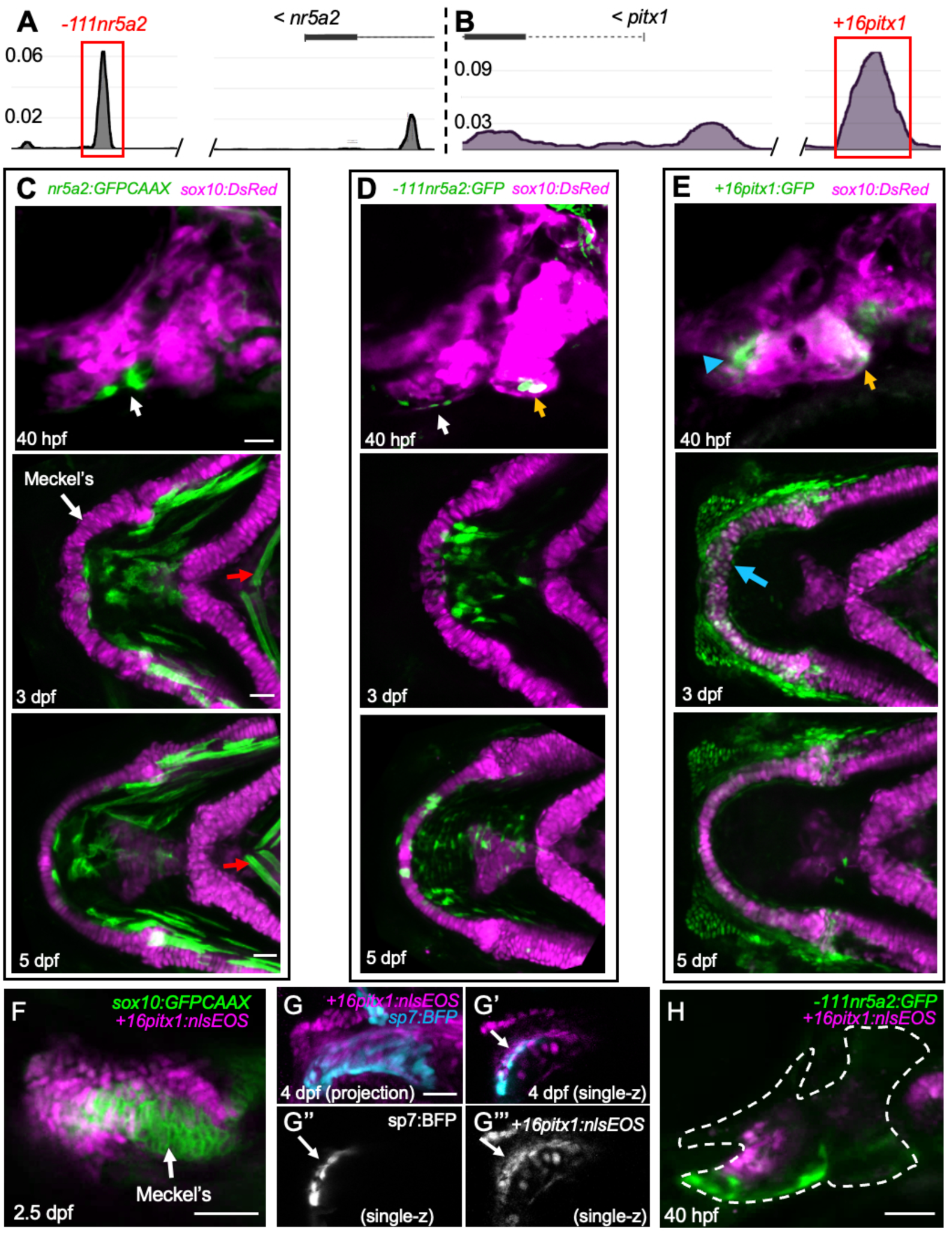
Identification of aboral *nr5a2* and oral *pitx1* enhancers. **(A,B)** Genome tracks for the *-111nr5a2 and +16pitx1* enhancers show the proportion of aggregated mesenchyme cells with indicated chromatin accessibility in CNCC snATAC-seq datasets from 36 hpf zebrafish (Fabian et al., 2022). **(C-E)** Activities of a *nr5a2:GFPCAAX* knock-in allele and *-111nr5a2:GFP* and *+16pitx1:nlsEOS* transgenes. *sox10:DsRed* labels CNCCs of the first two pharyngeal arches at 40 hpf (lateral view) and facial chondrocytes at 3 and 5 dpf (ventral view). Both *nr5a2:GFPCAAX* and *-111nr5a2:GFP* label aboral mandibular CNCCs (white arrows), and *+16pitx1:nlsEOS* labels oral mandibular CNCCs (blue arrowhead) at 40 hpf. At 3 and 5 dpf, *nr5a2:GFPCAAX* and *- 111nr5a2:GFP* label mesenchyme posterior to Meckel’s cartilage, and *+16pitx1:nlsEOS* labels mesenchyme anterior to Meckel’s cartilage and some chondrocytes at the distal tip of Meckel’s (blue arrow). The *nr5a2:GFPCAAX* knock-in allele also has activity in muscles (red arrows), while *-111nr5a2:GFP* and *+16pitx1:nlsEOS* are CNCC-specific but have additional activity in the second arch (yellow arrows) not reflective of endogenous *nr5a2* and *pitx1* expression. **(F, G)** Relative to the chondrocyte transgene *sox10:GFPCAAX* and osteoblast transgene *sp7:BFP*, *+16pitx1:nlsEOS* shows activity in ventral-most chondrocytes and surrounding mesenchyme of Meckel’s cartilage (F) and the osteoblasts and surrounding mesenchyme of the dentary bone (arrows, G). **(H)** At 40 hpf, *-111nr5a2:GFP* and *+16pitx1:nlsEOS* display complementary activity in the aboral and oral mandibular domains, respectively. The first two arches are outlined based on brightfield imaging (not shown).

We next examined regulation of aboral *-111nr5a2* and oral *+16pitx1* enhancers. Similar to regulation of endogenous *nr5a2* expression (**Figure 3**, **Figure 4**), we found that Bmp inhibition with Dorsomorphin eliminated *-111nr5a2:GFP* and *nr5a2:GFPCAAX* aboral activity, with Fgf inhibition with SU5402 expanding *- 111nr5a2:GFP* activity into the oral domain (**Figure 8A-F**). We also uncovered 6 sequences matching the consensus Hand2 E-box binding motif (CAnnTG) (**Figure 8G**), consistent with accessibility of -111nr5a2 overlapping with expression of the Bmp target and E-box factor *hand2* in our single-cell datasets (**Figure S2**). Consistent with Bmp-dependent Hand2 regulation of *nr5a2* (**Figure 6O**), mutation of all 6 motifs to AAnnTT severely reduced *-111nr5a2:GFP* activity in stable transgenic lines at 40 hpf and 6 dpf (**Figure 8J,M**). Conversely, mutation of two predicted SMAD binding motifs had no effect on *-111nr5a2:GFP* activity (**Figure 8I,L**).

**Figure 8.**
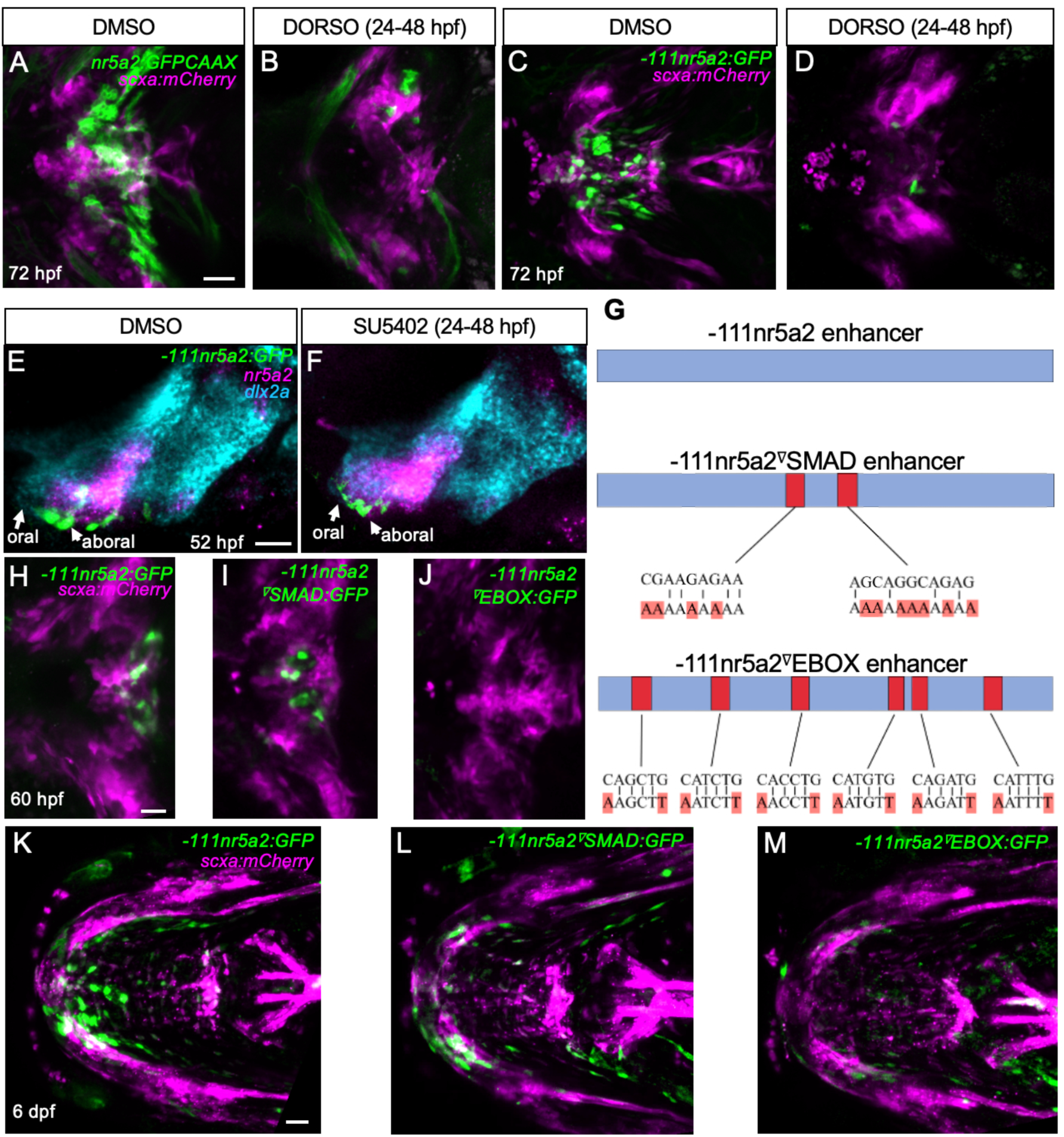
*111nr5a2* enhancer activity is regulated by Bmp and Fgf signaling and requires E-box motifs. **(A-D)** In ventral views of the lower jaw, *nr5a2:GFPCAAX* (A,B) and *- 111nr5a2:GFP* (C,D) show overlapping expression with the connective tissue marker *scxa:mCherry* in DMSO-treated controls. Treatment with the Bmp inhibitor Dorsomorphin (DORSO) from 24-48 hpf results in loss of CNCC transgene expression, while muscle expression remains in *nr5a2:GFPCAAX* embryos. **(E,F)** Anti-GFP staining shows activity of *-111nr5a2:GFP* relative to mRNA expression of *nr5a2* and the pan-CNCC marker *dlx2a* at 52 hpf. Compared to DMSO-treated controls, embryos treated with the Fgf inhibitor SU5402 from 24-48 hpf show expansion of *-111nr5a2:GFP* and *nr5a2* mRNA into the oral domain. **(G)** Schematic of the *-111nr5a2* enhancer and positions and nucleotide changes of the 2 SMAD and 6 E-box motifs separately mutated in transgenic assays. **(H-M)** Relative to the connective tissue marker *scxa:mCherry*, ventral views of the lower jaw show loss of activity at 60 hpf and reduced activity at 6 dpf when all 6 E-box motifs are mutated but not when the 2 SMAD motifs are mutated. Scale bars = 25 μM.

The +*16pitx1* DAR is conserved with a region near the mouse *Pitx1* gene, with both fish and mouse versions containing two conserved putative ETS binding motifs (GGAA) (**Figure 9A**). Chromatin accessibility of the enhancer also overlaps with expression of the Fgf8a-dependent oral mandibular ETS factor *erm* (*etv5b*) (Roehl and Nüsslein-Volhard, 2001) (Figure S2C). Consistently, SU5402-mediated Fgf inhibition eliminated and DMH1-mediated Bmp inhibition slightly expanded +*16pitx1:nlsEOS* activity at 40 hpf (**Figure 9B-D**). In addition, mutation of the predicted ETS sites to CCTT severely reduced *+16pitx1:nlsEOS* activity in stable transgenic lines at 40 hpf, 3 dpf, and 5 dpf (**Figure 9E-J**). These findings support Bmp-Hand2 and Fgf-ETS modules directly regulating aboral *nr5a2* and oral *pitx1* expression, respectively.

**Figure 9.**
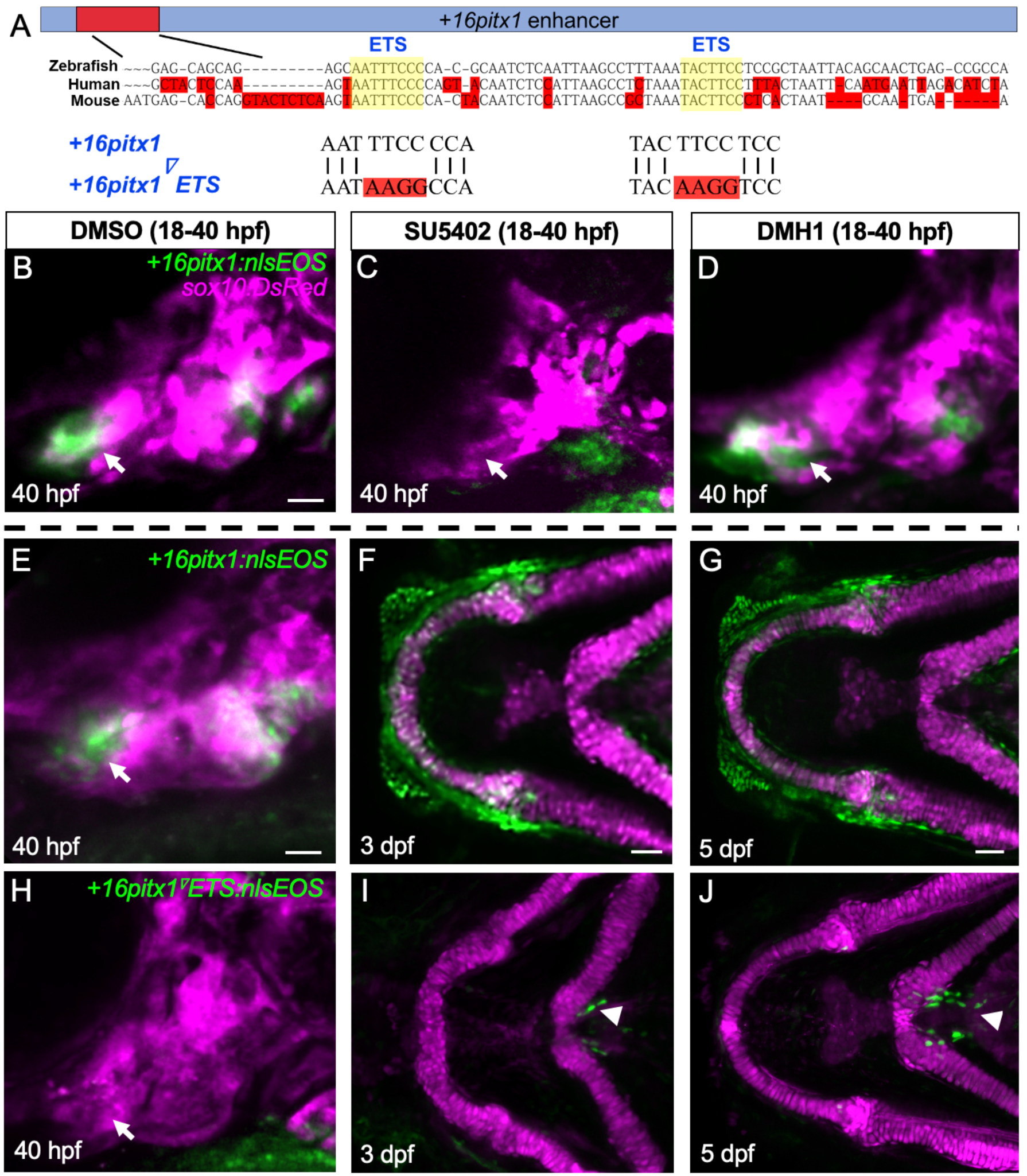
*+16pitx1* enhancer activity requires Fgf signaling and ETS motifs. **(A)** Schematic of the *+16pitx1* enhancer, position of the region conserved with mouse and humans and containing 2 ETS motifs, and nucleotide mutations used to destroy the ETS motifs. **(B-D)** Relative to DMSO-treated controls, oral mandibular expression of *+16pitx1:nlsEOS* (arrows) is lost upon treatment with the Fgf inhibitor SU5402 but not with the Bmp inhibitor DMH1 from 18-40 hpf. In these lateral views of the first two pharyngeal arches, *sox10:DsRed* labels CNCCs in magenta. **(E-J)** Relative to *sox10:DsRed* labeling of arch CNCCs at 40 hpf and jaw cartilages at 3 and 5 dpf, mutation of both ETS sites in the *+16pitx1* enhancer abolishes oral mandibular transgenic activity, although some expression remains in midline tendons attaching to the ceratohyal cartilages (arrowheads). Scale bars = 25 μM.

## DISCUSSION

Our findings reveal how Bmp signaling from the ventral ectoderm, Fgf signaling from the oral ectoderm, and Hh signaling from the medial endoderm interact to compartmentalize the mandibular arch into aboral, oral-lateral, and oral-medial domains that give rise to distinct jaw cell fates. Despite zebrafish lacking oral teeth, the roles of the these signaling pathways in mandibular arch patterning appear to be largely conserved with mammals, including the use of conserved regulatory sequences such as the *+16pitx1* oral enhancer.

In contrast to the oral domain, less was known about how the aboral domain of the mandibular arch is established. Prior work in mouse mandibular explants had defined *Lhx8* as a marker of the rostral domain and *Gsc* as a marker of the caudal domain (Tucker et al., 1999), which aligns with single-cell RNA sequencing studies of the mouse mandible (Xu et al., 2019b). Mandibular domains expressing *Foxf1* and *Foxf2* have also been described as oral (Xu et al., 2019b). We find that *foxf1* and *pitx1* have partially overlapping expression along the mediolateral axis of the oral domain, with co-expression of *gsc* and *nr5a2* in a non-overlapping domain posterior to *pitx1* and *foxf1*. *Gsc* and *Nr5a2* are also co-expressed in the caudal domain of the mouse mandibular arch, with *Gsc* mutants showing similar middle ear defects to conditional *Nr5a2* mutants (Chen et al., 2023; Yamada et al., 1995). We therefore define the *gsc*+; *nr5a2*+ domain as aboral and the *pitx1*+ and *foxf1*+ domains as oral-lateral and oral-medial regions, respectively.

Edn1 signaling appears to be broadly required for ventral mandibular gene expression in each sub-domain, as *gsc*, *nr5a2*, *pitx1*, and *foxf1* expression are all reduced or lost in zebrafish *edn1* mutants ((Askary et al., 2017; Talbot et al., 2010; Xu et al., 2018) and this study). In mouse, *Gsc* and *Pitx1* expression is similarly lost in *Endothelin-A receptor* mutants (Clouthier et al., 1998; Ozeki et al., 2004), and both *Gsc* and *Pitx1* expression are expanded into the maxillary domain upon Edn1 misexpression (Sato et al., 2008). In contrast, Fgf, Bmp, and Hh signaling have more restricted roles in mandibular sub-domain patterning. Our finding that Fgf signaling promotes oral *pitx1* and represses aboral *gsc* and *nr5a2* expression in zebrafish is consistent with mouse mandibular explant experiments in which exogenous FGF8 promotes oral *Pitx1*, *Pitx2*, *Lhx6*, and *Lhx8* expression and represses aboral *Gsc* expression (Grigoriou et al., 1998; St.Amand et al., 2000; Tucker et al., 1999). Conversely, we find that Bmp signaling is necessary and sufficient for aboral *gsc* and *nr5a2* expression. Whereas we did not observe expansion of *pitx1* expression upon Bmp inhibition, Bmp4 misexpression did repress oral *pitx1* expression, consistent with studies in mouse mandibular explants (Tucker et al., 1999). Whereas previous studies have shown Hh signaling to be necessary and sufficient for *Foxf1/foxf1* and *Foxf2/foxf2* expression in the medial mandibular domain (Xu et al., 2018; Xu et al., 2019a), we found that *foxf1* expression was not dramatically altered by inhibition of Fgf or Bmp signaling. These findings suggest a model in which *fgf8a* from the oral ectoderm and *shha* from the underlying endoderm act relatively locally to specify lateral and medial oral sub-domains, respectively, while *bmp4* from the ventral ectodermal margin specifies the aboral domain. However, these signaling pathways also likely interact in the mandibular arch as endodermal Shh induces oral ectodermal expression of both *Fgf8* and *Shh* (Brito et al., 2006; Haworth et al., 2007), Bmp4 represses oral ectodermal *Fgf8* expression (St.Amand et al., 2000), and mandibular Bmp signaling expands upon loss of Hh signaling (Xu et al., 2019a).

The analyses of transcription factor mutants and the sequence-conserved oral*+16pitx1* enhancer and aboral *-111nr5a2* enhancer shed light on how signaling pathways are interpreted to establish mandibular sub-domains. Previous studies have shown that the ETS transcription factors *etv4* (*pea3*) and *etv5b* (*erm*) have Fgf8a-dependent expression in the oral mandibular domain of zebrafish (Roehl and Nüsslein-Volhard, 2001), and we found that two predicted ETS sites conserved between fish and mouse are required for the oral activity of the *+16pitx1* enhancer. We also found that aboral *gsc* and *nr5a2* but not oral *pitx1* expression require the Hand2 bHLH transcription factor, consistent with previously reported specific loss of *gsc* expression in the ventral mandibular arch of zebrafish *hand2* mutants (Talbot et al., 2010), and partial reduction of *Gsc* in a mouse hypomorphic *Hand2* arch-specific enhancer deletion allele (Funato et al., 2016). Whereas both Edn1 and Bmp signaling are required for *Hand2/hand2* expression (Charité et al., 2001; Yanagisawa et al., 2003), we had previously shown that *hand2* expression was more strongly induced by Bmp4 than Edn1 in zebrafish (Zuniga et al., 2011). Consistent with Hand2 mediating Bmp4 induction of aboral gene expression, we found that expansion of *nr5a2* and *gsc* expression upon Bmp4 misexpression was abrogated by *hand2* loss. Moreover, mutation of six bHLH motifs in the *-111nr5a2* enhancer severely reduced its activity, although we cannot rule out that other bHLH factors such as Twist1 could bind these elements. While we did not address how Fgf signaling represses *gsc* and *nr5a2* expression, a previous study provided correlative evidence that the Fgf8-dependent oral factor Lhx8 restricts *Gsc* aborally (Tucker et al., 1999). In contrast, oral *Lhx8* expression was shown not to depend on *Gsc* function in mouse (Tucker et al., 1999), and we found no evidence in zebrafish that Gsc, Pitx1, and Nr5a2 cross-regulate their expression ((Chen et al., 2023) and this study). These findings support a model in which Fgf8 induces oral gene expression through ETS factors and Bmp4 induces aboral gene expression through Hand2, although there are likely other transcriptional inputs yet to be discovered.

Our findings also illuminate how compartmentalization of the ventral mandibular arch may specify distinct cell fates and sculpt the lower jaw. Photoconversion-based lineage tracing of the aboral *nr5a2*+/*gsc*+ domain showed contribution to ligament and stromal tissues but not cartilage or bone, consistent with our previous work demonstrating roles for Nr5a2 in promoting tendon, ligament, and salivary gland mesenchyme at the expense of cartilage fates in fish and mouse (Chen et al., 2023). In contrast, photoconverted oral and oral-aboral intermediate regions, which overlap with *pitx1*+ and *foxf1*+ oral domains, gave rise to bone and cartilage. While both the Fgf-dependent *pitx1*+ and Hh-dependent *foxf1*+ domains are non-overlapping with the Bmp-dependent *gsc+*; *nr5a2*+ aboral domain, *pitx1* and *foxf1* expression overlap with *pitx1*-only cells more laterally and *foxf1*-only cells more medially. The Shh-producing medial endoderm has been shown to be critical for Meckel’s cartilage formation (Eberhart et al., 2006; Lan and Jiang, 2009; Schwend and Ahlgren, 2009; Xu et al., 2019a), with ectopic production of Shh resulting in branched or even supernumerary Meckel’s cartilage (Balczerski et al., 2012; Xu et al., 2018). Shh may induce Meckel’s cartilage in part through Foxf1/2 genes as their loss in zebrafish and mouse result in truncation of Meckel’s cartilage, with the related Foxc1a/b factors acting to prime CNCCs for facial cartilage formation in zebrafish (Xu et al., 2018; Xu et al., 2019a). Moreover, CNCC-specific loss of *Fox1/2* or the Shh receptor *Smoothened* in mouse results in ectopic bone formation in the oral domain (Xu et al., 2019a), with Foxf1 misexpression in zebrafish inhibiting intramembranous bone formation (Xu et al., 2018). Shh from the medial endoderm also functions to induce *Shh* and *Fgf8* expression in the oral ectoderm (Brito et al., 2006; Haworth et al., 2007), with loss of *Shh* function in the mouse oropharyngeal epithelium resulting in complete absence of Meckel’s cartilage (Billmyre and Klingensmith, 2015). Similar to loss of *Foxf1/2*, deletion of *Fgf8* from the oral ectoderm (Trumpp et al., 1999) or *Pitx1* loss (Mitsiadis and Drouin, 2008) results in truncation of the lower jaw. Our analysis of the zebrafish *+16pitx1* enhancer that is conserved in mouse revealed that *pitx1*-expressing CNCCs give rise to the dentary bone and the distal region of Meckel’s cartilage. One possibility then is that integration of Shh and Fgf8 signaling would specify Meckel’s cartilage in *foxf1*+; *pitx1*+ CNCCs adjacent to the medial endoderm, with the more lateral *pitx1*-only region of the oral domain contributing to intramembranous bone and oral teeth (in some species), in addition to acting as a growth zone for distal extension of the mandible. Conversion of a *foxf1*+; *pitx1*+ domain to a *pitx1*-only domain in the absence of Hh signaling or Foxf1/2 genes might also explain the ectopic bone phenotypes observed in mice (Xu et al., 2019a). Future experiments will be needed to elucidate how Hh- and Fgf-dependent transcription factors interact to regulate multilineage differentiation and cell proliferation along the mediolateral axis of the oral domain to elongate and pattern the lower jaw.

## Materials and Methods

### Experimental model and subject details

All experiments on zebrafish (*Danio rerio*) were approved by the Institutional Animal Care and Use Committee of the University of Southern California (IACUC protocols #20771). Embryos and animals were humanely euthanized for experiments. Zebrafish were raised in vivarium under standard conditions at 28.5°C with health and water conditions monitored daily. The following zebrafish lines were used in this study: *Tg(sox10:EGFP-CAAX)^el375^, Tg(sox10:DsRed)^el10^, Tg(nr5a2:GFP-CAAX)^el874^,Tg(scxa:mCherry)^fb301^,Tg(sox10:kikGR)^el2^, Tg(+16pitx1_p1-Mmu.E1b:nlsEOS,cryaa:Cerulean)^el1056^,Tg(+16pitx1_p1noets-Mmu.E1b:NLSEOS,cryaa:Cerulean)^el1057^,Tg(-111nr5a2-Mmu.E1b:GFP,cryaa:Cerulean)^el880^,Tg(-111nr5a2noebox-Mmu.E1b:GFP,cryaa:Cerulean)^el1054^,Tg(-111nr5a2nosmad-Mmu.E1b:GFP,cryaa:Cerulean)^el1055^,Tg(-1.5hsp70l:Gal4)^kca4^, Tg(UAS:Bmp4,myl7:EGFP)^el49^,Tg(GBS-ptch2:EGFP)^umz23tg^, Tg(dusp6:d2EGFP)^pt6tg^, pitx1^el1058^, gsc^x59^, moz^b719^,* and *edn1^tf216b^.* Heterozygous mutant adult fish were raised in similar conditions and showed no visible defects, with homozygous mutant embryos produced by breeding for experimental analysis. All heat-shock inductions were performed by transferring embryos from 28°C to pre-warmed 38°C embryo media for 20 minutes. Individual identifiers for zebrafish lines are detailed in Key Resource Table 1.

### Generation of transgenic and mutant zebrafish lines

The *pitx1* mutant was generated via CRISPR-Cas9 targeted mutagenesis using the Alt-R system (Integrated DNA Technologies). We generated a full locus deletion with all protein-coding exons removed. Synthetic crRNAs were designed to target upstream the first exon, and downstream of the last exon (see Key Resource Table 1 for sequences). Standard Alt-R™ Cas9 nuclease was purchased from IDT. Lyophilized Alt-R crRNA and tracrRNA were resuspended in Nuclease-Free IDTE Buffer to a final concentration of 100 µM. The guide RNA duplex was assembled at a final concentration of 3 µM. The mixture was incubated at 95°C for 5 minutes, then cooled to room temperature on the benchtop. Alt-R™ Cas9 nuclease was diluted to a working concentration of 0.5 µg/µL in Cas9 working buffer (20 mM HEPES, 150 mM KCl, pH 7.5). The final ribonucleoprotein (RNP) complex for each guide was prepared by combining 3 µL gRNA duplex with 3 µL of the 0.5 µg/µL diluted Cas9 protein. This mixture was incubated at 37°C for 10 minutes to allow RNP complex formation, then cooled to room temperature. This mixture was microinjected into single-cell Tubingen zebrafish embryos (2 nL), and stable mutant carriers were identified by PCR of their embryos. A more detailed characterization of the mutant will be described elsewhere.

To generate transgenic lines, donor plasmids (final concentration of 20 ng/μl) were co-injected with Tol2 transposase RNA (final concentration of 25 ng/μl) into one-cell-stage embryos. Wild-type and modified enhancers for *-111nr5a2* (GRCz11:Chr22:22609540-22610198) and *+16pitx1* (GRCz11:Chr21: 45831534-4583219) were synthesized as gBlocks (Integrated DNA Technologies) and cloned using In-Fusion (Takara) in front of a *Mmu.E1b* minimal promoter, followed by either GFP or nlsEOS. Plasmids also contained *cryaa:Cerulean* as a co-selection marker driving lens fluorescence, flanked by Tol2 transposase sequences for integration. Three independent founder lines were identified for each construct, and consistent GFP/EOS expression patterns were observed in at least three embryos of each allele. A representative transgenic founder for each pattern was given an allele number, and its progeny used for all images presented in the manuscript. At least n≥6 individuals were analyzed for comparative transgenic activity experiments in this study.

### Reanalysis of zebrafish CNCC scRNAseq and snATACseq datasets

To visualize genomic accessibility near genes of interest, we utilized a preexisting cloupe (10x Genomics Cell Ranger) output file generated from isolated 1.5 dpf *sox10:Cre; actab2:loxP-BFP-STOP-loxP-DsRed* sorted CNCCs (Fabian et al., 2022). Individual peaks of interest were manually discovered by utilizing the peak view function in Loupe Browser 9.0.0 (10x Genomics), and candidate DNA sequences were referenced from the GRCz11 genome assembly annotated on Ensembl genome browser 115 (Ensembl). Sequences of interest were used to generate gBlocks (Integrated DNA Technologies) for cloning into transgenic expression plasmids as detailed above.

To generate feature plots to visualize enhancer accessibility and RNA expression patterns in individual cells, we utilized a previously generated dataset of CNCCs from 1.5 dpf fish heads (Fabian et al., 2022). Reanalysis was performed on the Signac/Seurat RDS object for the 1.5 dpf integrated snATACseq/snRNAseq dataset, which had been processed using the snapATAC bioinformatics pipeline, available at FaceBase under the RID “D-NKM4” (Fabian et al., 2022). Accessibility for peaks -111nr5a2 and +16pitx1 were visualized using the Featureplot function in Seurat (V5.2.0). To compare gene expression patterns and activity for the - 111nr5a2 and +16pitx1 enhancers, we utilized the function FeaturePlot with the blend = TRUE option. Cells positive for individual enhancer accessibility and individual gene expression patterns were co-visualized as UMAPs in Seurat (V5.2.0).

### In situ hybridization and whole-mount immunofluorescence

Embryos were grown to the indicated developmental stages and fixed in 4% PFA in PBS solution overnight at 4°C (Sigma). Fixed embryos were then incubated with 10 μg/ml of ProK diluted in PBST for 30 min. This was followed by three 5-min washes with PBST, then fixation in 4% PFA for 20 min. In situ hybridizations were performed using Molecular Instruments HCR (V3.0) technology following whole-mount zebrafish protocols (Choi et al., 2014; Trivedi et al., 2018). RNA HCR hairpins were synthesized by Molecular Instruments and detailed in Key Resource Table 1. After completion of the whole-mount protocol, samples were incubated for 15 min with PBDTx (5 ml 10x PBS, 0.5 g BSA, 500 ul DMSO, three drops of 1M NaOH, 250 ul of 20% TritonX solution in a total volume of 50 ml), followed by a 3-h blocking step in PBDTx with 2% goat serum (PBDT-GS) at room temperature. The primary antibody chicken anti-GFP (1:200, Abcam ab 13970) was diluted in PBDT-GS, incubated overnight at 4°C, followed by three 20-min washes with PBDTx. Secondary antibody Alexa Fluor 488 goat anti-chicken (1:300, Thermo Fisher A-11039) was diluted with PBDT-GS and incubated at room temperature for 5 h in the dark. Fish were then washed three times for 5 min each with PBS containing 1% TritonX and stored at 4°C in the dark in PBS until imaged.

To generate frontal sections of whole-mount in situ embryos, individuals were initially imaged sagittally and then transferred to a polystyrene 100mmx20mm petri dish filled with roughly 50 mL of 5x SSCT (750 mM NaCl, 75 mM trisodium citrate with 0.1% Tween-20, pH 7.0). Individual heads were dissected in a frontal manner, immediately posterior to the eye, with a 22.5° cutting angle microknife (Fine Science Tools). Dissected heads were transferred from the petri dish and oriented in a frontal manner in warmed (42°C) liquid 0.2% agarose in PBS in a 35-mm glass-bottom dish with uncoated #1.5 coverslip and 20 mm glass diameter (MatTek). Once agarose solidified, embryos were then imaged using confocal microscopy.

### Imaging

Images of live and fixed zebrafish, as well as individual sections, were captured on a Zeiss LSM800 confocal microscope using the ZEN Blue software with 10× or 20× objectives. Images were taken as three-dimensional stacks using the z-stack function and displayed as maximum intensity projections or individual slices processed with Fiji software (ImageJ). Images were adjusted for brightness in Fiji software, with equal processing within experiments. Photoconversion experiments for *sox10:kikGR* and *+16pitx1:nlsEOS* were done using the area of interest function to select regions of interest for 405 nm laser exposure. For *sox10:kikGR* photoconversion experiments, each indicated region was targeted using the area of interest function followed by 75% 405 nm laser exposure for 2 min. For the *+16pitx1:nlsEOS* lineage tracing experiments, the oral region was selected and exposed by 75% 405 nm laser power for 2 min. Photoconversion was confirmed by brief 546 nm laser exposure, and the embryos were then transferred to embryo media in the dark until the indicated timepoint.

### Statistics

Except where indicated, n≥6 individuals were analyzed for reported expression and transgenic activity patterns.

## Supporting information

Key Resource Table 1

## Acknowledgements

We thank Megan Matsutani and Maya Lujan for fish care, and Kuang-Tse Wang for helpful comments on the manuscript. Funding was provided by National Institute of Dental and Craniofacial Research grants R35DE027550 and R21 DE029656 to J.G.C., and T90DE021982 and F32DE033911 to E.P.

## Author contributions

E.P.- Conceptualization, Data curation, Formal analysis, Investigation, Methodology, Validation, Visualization, Writing the manuscript. M.J.- Resources. L.B.- Investigation. H.C.- Investigation, Data curation, Formal analysis J.G.C.-Conceptualization, Formal analysis, Funding acquisition, Investigation, Methodology, Project administration, Supervision, Visualization, Writing – review and editing.

## Competing interests

The authors declare no competing interests.

---

**Figure S1.**
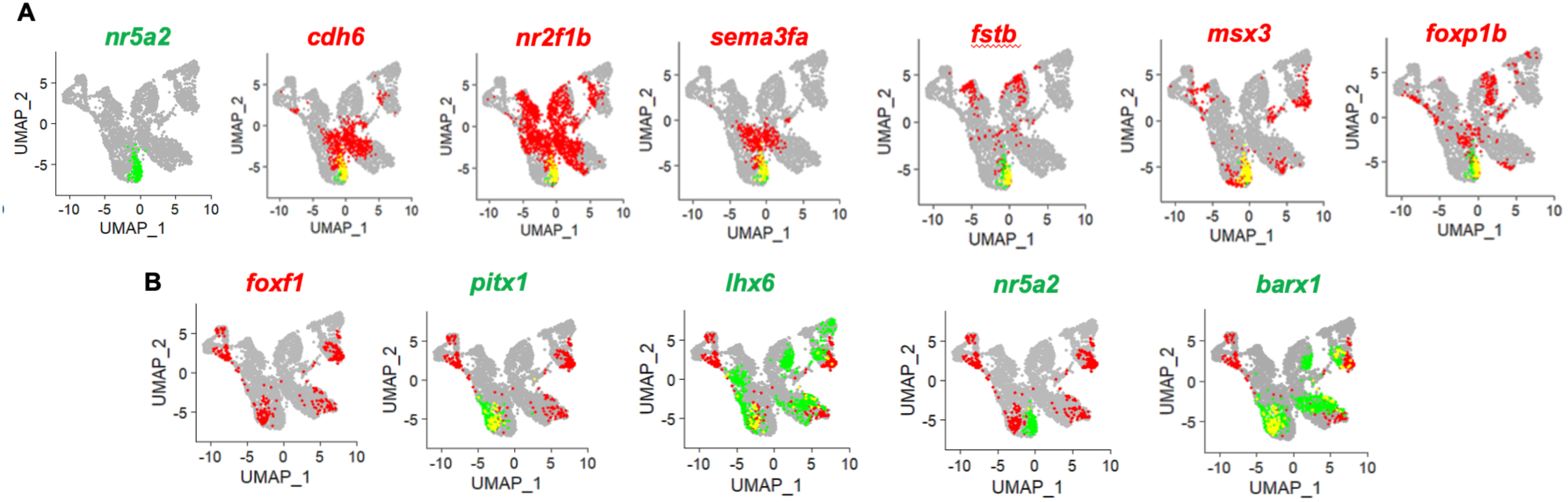
Predicted oral-aboral gene activity from snapATAC datasets of CNCCs from 36 hpf zebrafish. **(A)** Feature plots show predicted mRNA expression of agoral genes (red) relative to activity of *nr5a2* mRNA (green). **(B)** Feature plots show predicted mRNA expression of oral *pitx1* and *lhx6*, aboral *nr5a2*, and pre-chondrogenic *barx1* (green) relative to *foxf1* (red).

**Figure S2.**
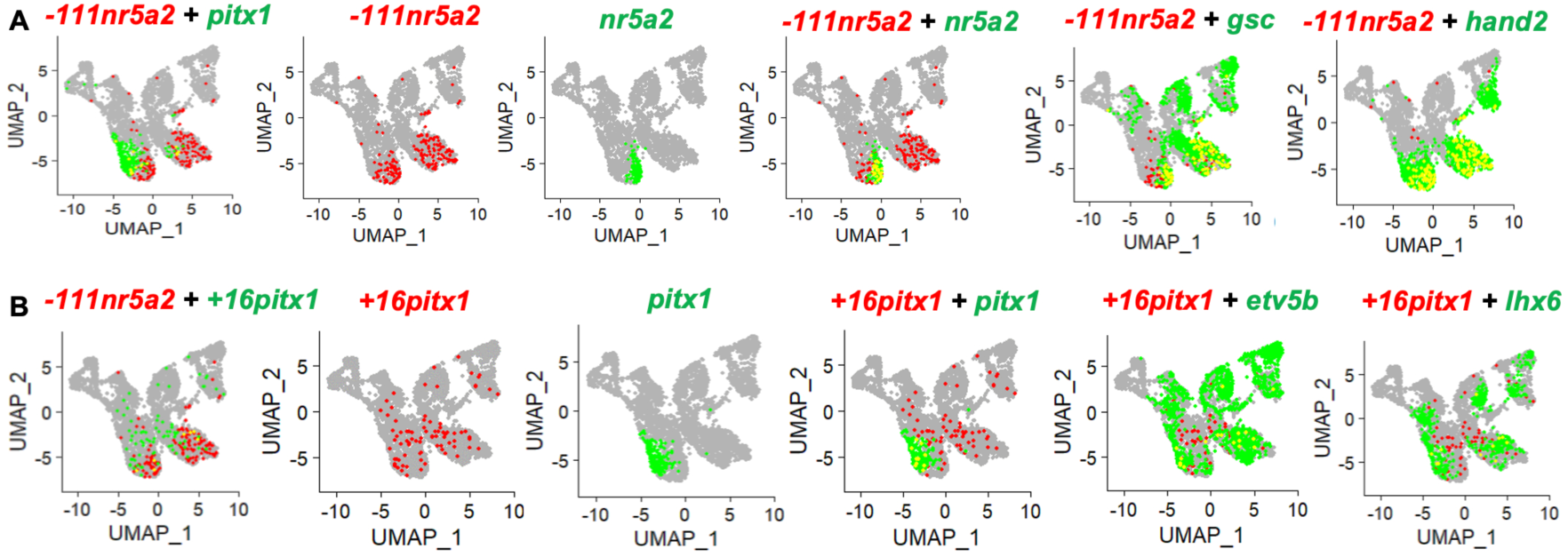
Analysis of relative chromatin accessibility for *-111nr5a2* and *+16pitx1* enhancers. **(A)** Feature plots for *-111nr5a2* enhancer accessibility (red) relative to select mRNA expression. **(B)** Feature plots show *+16pitx1* enhancer accessibility relative to the *-111nr5a2* enhancer and select mRNAs.

## Notes

### Competing Interest Statement

The authors have declared no competing interest.

